# Yeast Securin is trafficked through the endomembrane system to regulate cell cycle arrest during the DNA damage response

**DOI:** 10.1101/2019.12.15.876896

**Authors:** Brenda R. Lemos, David P. Waterman, James E. Haber

**Affiliations:** Department of Biology and Rosenstiel Basic Medical Sciences Research Center, Brandeis University, Waltham MA 02454-9110

## Abstract

The yeast securin protein, Pds1, belongs to a class of highly conserved eukaryotic proteins that regulate the timing of chromatid segregation during mitosis by inhibiting separase, Esp1. During the metaphase to anaphase transition Pds1 is degraded by the ubiquitin-proteasome system in the nucleus, unleashing Esp1’s protease activity to cleave cohesin surrounding sister chromatids. In response to DNA damage Pds1 is phosphorylated and stabilized, stalling mitotic progression to preserve genomic integrity. In addition, during the DNA damage checkpoint response, securin and separase are partially localized in the vacuole. Here we genetically dissect the requirements for securin’s vacuolar localization and find that it is dependent on the Alkaline Phosphatase (ALP) endosomal transport pathway but not autophagy. Blocking retrograde traffic between the Golgi and the endoplasmic reticulum, by inhibiting COPI vesicle transport, drives Pds1 into the vacuole, whereas inhibiting antegrade transport by disrupting COPII-mediated traffic, results in the unexpected loss of Pds1 and the extinction of DNA damage-induced cell cycle arrest. We report that the induction of ER stress, either genetically or by treating cells with dithiothreitol, is sufficient to extinguish the DNA damage checkpoint, suggesting crosstalk between these two pathways. These data highlight new ways in which Pds1 and Esp1 are regulated during a DNA damage induced G2/M arrest and its requirement in the maintenance of the DNA damage checkpoint response.

## Introduction

A single irreparable double strand break (DSB) activates the DNA damage checkpoint response in budding yeast. The DNA damage response is mediated by two PI3K-like kinases, Mec1 and Tel1 (ATR and ATM in mammals) that in turn phosphorylate several transducer and effector proteins, all to ensure G2/M arrest. Arrest prior to anaphase prevents cells from completing mitosis with broken chromosomes, allowing time for repair to take place (1, 2). Failure to activate the DNA damage checkpoint response can result in genomic instability (3) and in higher eukaryotes may lead to disease (4).

In *Saccharomyces cerevisiae*, Pds1(securin) binds to Esp1(separase) during anaphase onset and together are trafficked to the nucleus where the Anaphase Promoting Complex (APC), along with its cofactor Cdc20, ubiquitinates Pds1, which is then destroyed by the ubiquitin-proteasome system (5–8). Degradation of Pds1 allows Esp1 to cleave the Scc1 subunit of cohesin, allowing sister chromosome separation and the completion of mitosis (9, 10). In the presence of even a single DSB the DNA damage response is activated; Pds1 is phosphorylated and stabilized by the Chk1 kinase, which prevents Pds1’s ubiquitination and thus blocks Esp1’s protease activity and maintains G2/M arrest (11–13).

Cells suffering a single irreparable DSB arrest for 12-15 h, after which they turn off the DNA damage checkpoint and re-enter mitosis in a process termed adaptation (14–17). Several adaptation-defective mutations have been identified that render cells incapable of adapting and lead to permanent G2/M arrest (2). Why cells undergo adaptation in the presence of DNA damage is unknown, but this phenomenon is not limited to budding yeast. Indeed, adaptation has been described in both *Xenopus* egg extracts and in mammalian cells (18, 19). Adaptation thus allows cells with unrepaired DNA damage to divide and enhances the risk of genome instability. Therefore, it is of general interest to determine what mutations and conditions renders cells adaptation-defective when faced with irreparable DNA damage.

One set of adaptation-defective mutants that have been previously characterized are the deletions of Vps51 and other proteins belonging to the Golgi-associated retrograde protein complex (GARP) (14). Whereas Pds1-GFP normally localizes to the nucleus during DNA damage-induced G2/M arrest, in *vps51∆* cells Pds1-GFP instead largely forms puncta in the cytoplasm. Both the localization and adaptation defects were rescued by deleting the vacuolar protease Pep4 (14), which is also required for the maturation of other resident vacuolar proteases (20, 21). These observations along with the identification of a DNA damage-induced autophagy pathway termed, Genotoxin-induced Autophagy (GTA) (22), led us to suspect that Pds1 was a target of GTA. Here we show that Pds1 is targeted to the vacuole independently of autophagy and instead requires components of the alkaline phosphatase transport pathway (ALP) (23). Inhibiting Pds1’s vacuolar localization shortens cell arrest after the DNA damage checkpoint response is triggered, causing cells to resume mitosis more quickly in the presence of an unrepaired DSB. Additionally, we find a role for the major nuclear exporter Crm1 (Xpo1) in exporting Pds1 during the DNA damage checkpoint response and during valproic acid (VPA)-induced nuclear delocalization.

The dynamic movements of Separase and Securin are evident by interfering with vesicle transport. When COPI vesicles are inhibited and retrograde traffic is blocked, Pds1 is more strongly driven into the vacuole. Conversely, upon conditional inhibition of COPII vesicles to block antegrade traffic, Pds1 is unexpectedly degraded and the DNA damage checkpoint response is turned off. This result is reproduced by treatment of cells with dithiothreotol (DTT) that also induces ER stress (24). These results suggest that induction of ER stress takes precedence over the DNA damage response. Taken together our results indicate that Pds1 is highly dynamic during DNA damage induced G2/M arrest, offering insights into a key regulator of the DNA damage checkpoint response.

## Results

### Pds1 Localizes to the vacuole independently of autophagy

We have previously characterized an Mec1- and Tel1-dependent pathway for DNA damage-induced autophagy (22). We speculated that Pds1 is a target of GTA since we had shown Pds1 localized to vacuoles following a DSB. Since globular GFP is mostly resistant to vacuolar proteases, targets of autophagy can be monitored by fusing GFP to proteins and monitoring the appearance of GFP in the vacuole (25–27). However, GFP-tagged Pds1 in wild type cells was only seen localizing to the nucleus during DNA damage induced G2/M arrest but when the vacuolar protease, *PEP4*, was deleted, Pds1-GFP was visible in the vacuole (14) (Fig 1A). *PEP4* encodes for proteinase A that is required for the activation of other vacuolar proteases proteinase B and C, (28, 29). In the absence of Pep4, protein degradation is reduced by 20-30%, causing an accumulation of proteins in the vacuole and allowing visualization of GFP-tagged proteins otherwise not visible (30). Under these conditions Esp1-mCherry (Separase) also localizes to vacuoles, colocalizing with Pds1-GFP (Fig. 1B). This vacuolar localization allows us to study the genetic requirements for Pds1’s vacuolar degradation.

**Figure 1:**
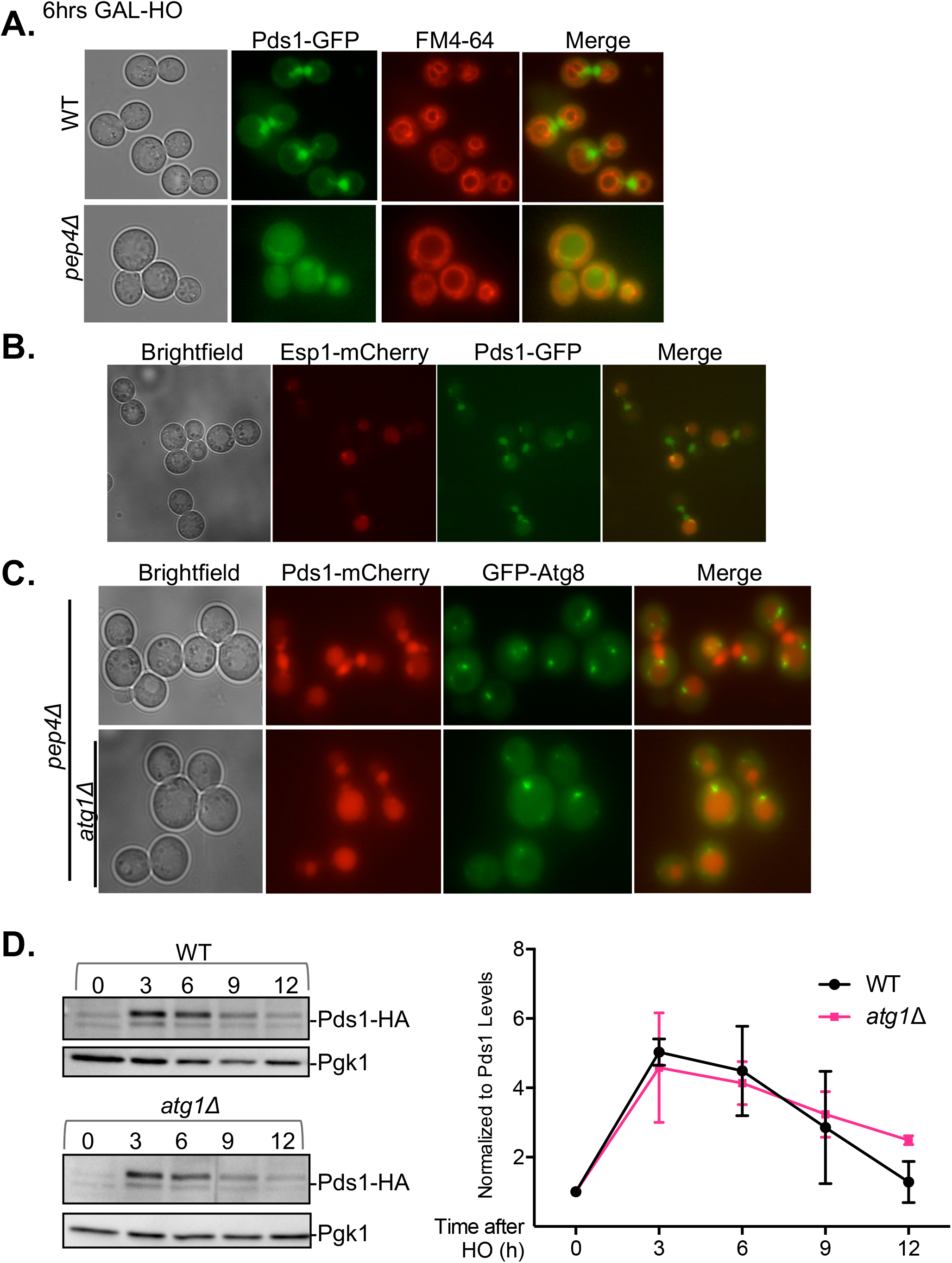
Pds1 localizes to vacuoles independently of autophagy A) Pds1-GFP localization in cells experiencing a single HO induced DSB. In the absence of the vacuolar protease Pep4, Pds1 localizes to the vacuole. FM4-64 highlights the vacuolar membrane. B) Esp1-mCherry co-localizes with Pds1-GFP in the vacuole. C) Pds1-mCherry localization in the absence of the core autophagy kinase Atg1. D) Quantification of Pds1 levels in cells experiencing a single HO-induced DSB in JKM179 background and in *atg1∆* cells.

Reasoning that autophagy is a likely pathway by which Pds1 is delivered to the vacuole during the DNA damage response, we deleted core autophagy proteins Atg1 and Atg13, which are both required for autophagic induction in yeast <ref>, in the *pep4*∆ background and monitored Pds1-mCherry localization 6 h after HO induction. Surprisingly, Pds1 still localized to the vacuole in the absence of these core autophagy proteins (Fig. 1C). Additionally, we monitored Pds1 abundance during the DNA damage response and observed a 5-fold increase in Pds1 protein levels, peaking at 3 h after HO induction and falling back to initial levels by 12 h (Fig. 1D). This profile was unchanged in *atg1*∆ cells (Fig. 1D). We therefore conclude that Pds1’s vacuolar localization does not require autophagy.

### Pds1 vacuolar localization requires the ALP pathway

We next extended our analysis to encompass other vacuolar-targeting pathways. The CPY pathway delivers vacuolar hydrolases and proteins from the Golgi to the vacuole through a pre-endosomal/pre-vacuolar compartment (PVC). The ALP pathway (alkaline phosphatase) delivers proteins directly from the Golgi to the vacuole. To test whether Pds1 is delivered to the vacuole via the CPY pathway, we monitored its localization in a *pep12*∆ mutant, which lacks a t-SNARE involved in vesicle fusion with the late endosome (31). Pds1 vacuolar localization persisted in *pep4*∆ cells lacking without Pep12, as judged by overlapping GFP signal with the vacuolar lumen stain CMAC (Fig. 2D). This was also the case in cells lacking the sorting receptor Pep1 that is also required for the CPY pathway (Fig. 2D) (32). Thus, Pds1 is not delivered to the vacuole via the CPY pathway.

**Figure 2:**
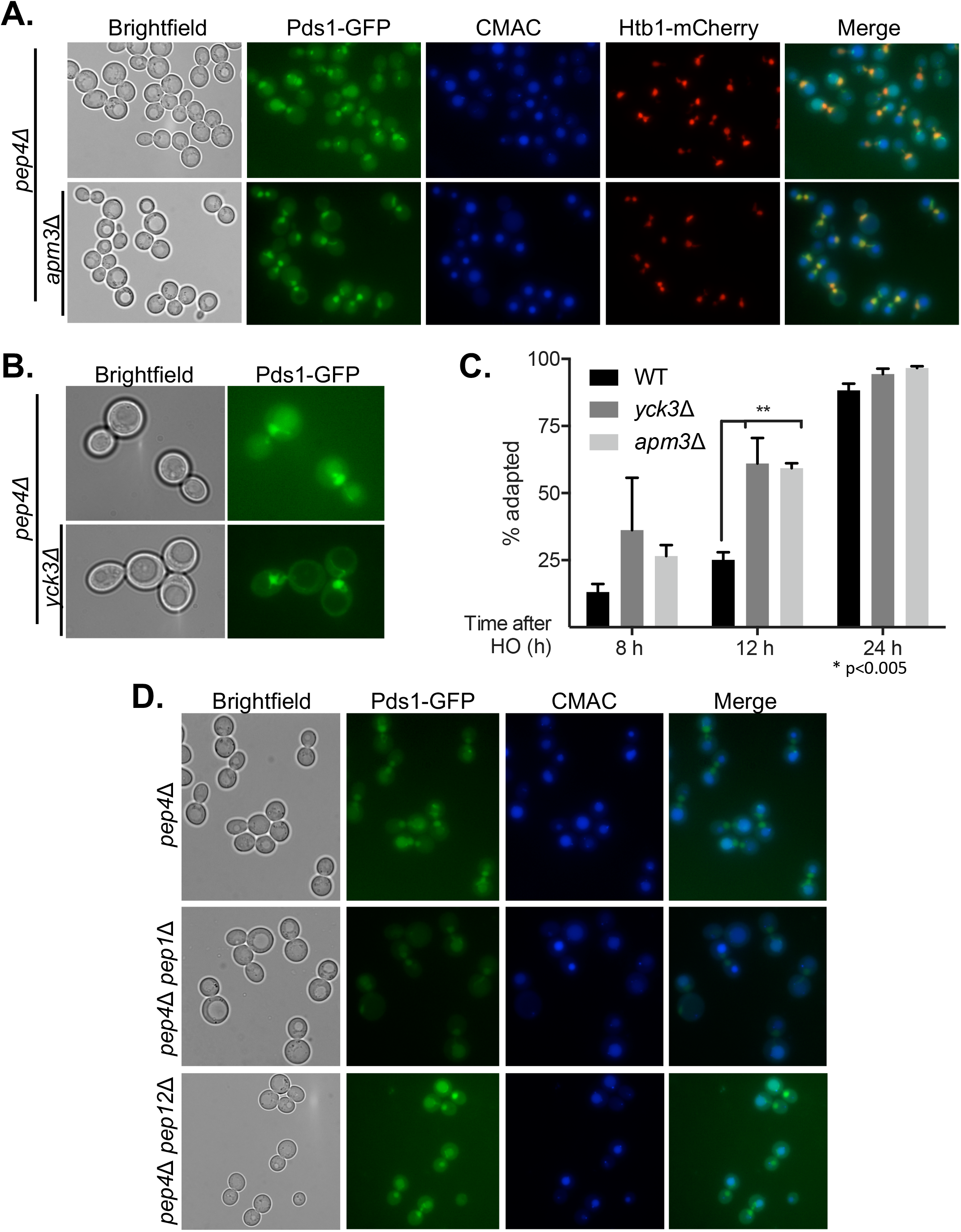
Pds1 localizes to vacuoles using components of the ALP pathway A) Pds1-GFP localizes to vacuoles in *pep4∆* but localization is blocked in *pep4∆ apm3∆* cells b) In the absence of the vacuolar kinase Yck3, Pds1-GFP does not localize to the vacuole. B) 24 h adaptation assay in WT, *yck3∆* and *apm3∆* cells. *p<0.005 students t-test with Bonferroni correction from three separate experiments. C) Pds1-GFP localization in the absence of components of the CPY pathway, *pep12∆* and *pep1∆*.

We then examined the ALP pathway. The ALP pathway requires the heterotetrametric AP-3 adaptor complex to deliver vesicles from the Golgi to the vacuole (33). We monitored Pds1’s localization in *apm3*∆ cells, which lack the µ3a subunit of AP-3 (34), after induction of a single HO-mediated DSB and found that Pds1 no longer localized to the vacuole in this mutant (Fig. 2A). To substantiate this finding we deleted the vacuolar kinase Yck3 required for the fusion of ALP vesicles with the vacuole (35). Pds1’s vacuolar localization was also blocked in *yck3*∆ cells (Fig. 2B). Taken together, these results suggest that Pds1 localizes to the vacuole using components of the ALP pathway.

### apm3∆ and yck3∆ cells adapt faster after DNA damage

We then asked how the ALP mutations would affect cell cycle arrest after DNA damage, in particular how cells would arrest and adapt after induction of a single unrepaired DSB. Cells were grown overnight in YEP supplemented with 3% lactic acid and plated on YEP plates containing 2% galactose, which induces a single DSB created by HO endonuclease (17). G1 cells were micromanipulated and monitored for 24 h to determine the time at which cells adapted. WT cells arrest at G2/M for 12-15 h then turn off the DNA damage checkpoint and resume mitosis (17) (Fig. 2C). Although only 25% of WT cells had adapted by 12h, over 50% of *apm3*∆ and *yck3*∆ cells had adapted (Fig. 2C). These data suggest that Pds1’s vacuolar localization is required for proper maintenance of G2/M arrest after DNA damage.

### Pds1 localizes to the vacuole in the absence of COPI vesicles

Given that the ALP pathway in responsible for Pds1’s localization to the vacuole during the DNA damage response, we reasoned that perturbing other cell endomembrane trafficking components may alter Pds1’s localization. To this end we examined the role of COPI vesicles, which are a coatomer proteins that deliver vesicles from the *cis*-Golgi to the ER, with the rationale that if Pds1 localizes to the vacuole during the DNA damage response, it may also be trafficked back to the nucleus via retrograde trafficking (36). Therefore, if retrograde trafficking is blocked, Pds1 might accumulate in the vacuole. We tagged the delta subunit of COPI vesicles, Ret2 (37), with an auxin-inducible degron (AID) (38, 39). Incorporation of AID at the C-terminus of Ret2 and degradation of the Ret2 subunit upon auxin treatment was confirmed via western blot (Fig. S1). Cultures were grown overnight to exponential phase and galactose was added to induce a DSB for 2 h prior to auxin treatment. Fewer than 25% of cells had Pds1 in the vacuole before IAA treatment whereas over 75% of cells had Pds1 in the vacuole in the absence of COPI vesicles, even in the presence of Pep4 (Fig. 3A). These results suggest that in the absence of COPI vesicles, Pds1 is cycled into the vacuole and fails to be retrograde transported back toward the nucleus.

**Figure 3:**
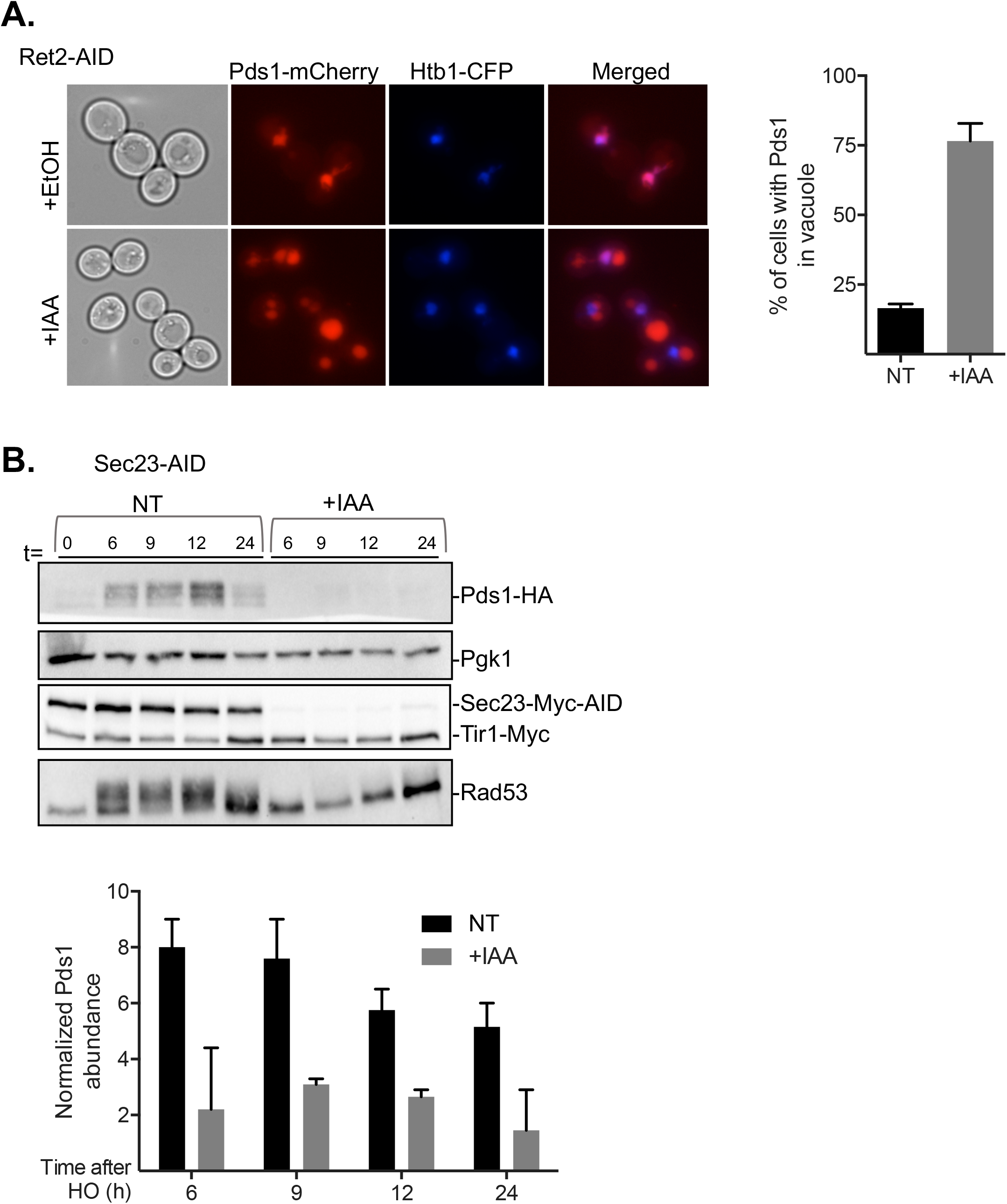
Pds1’s localization and concentration in the absence of COPI and COPII vesicles A) Western blot showing degradation of COPI subunit Ret2-AID-MYC upon IAA treatment. Pds1-mCherry localization in the background of Ret2-AID with quantification of the fraction of cells with Pds1 in the vacuole before and after IAA treatment. B) Western blot and quantification (C) of Pds1 in the absence of the COPII subunit Sec23.

### Inhibition of COPII vesicles leads to degradation of Pds1 and inactivation of the DNA damage checkpoint response

Since Pds1 travels to the vacuole via the ALP pathway, we presumed that it undergoes anterograde traffic from the ER to the Golgi and therefore might also depend on COPII vesicle transport. We tested whether conditionally inhibiting COPII vesicles via Sec23-AID depletion altered Pds1’s vacuolar localization. Cells were grown overnight in YEP-Lac to exponential phase and treated with 2% galactose to induce a DSB. 2 h after DSB induction, auxin was added to cultures to deplete Sec23. In *pep4*∆ cells treated only with galactose, Pds1 localized to both the vacuole and nucleus. As expected, in cells only treated with galactose to induce a DSB, Pds1 accumulated in the cells during arrest, with protein concentration peaking around 6 h after induction of a DSB and falling thereafter (Fig. 3B). Surprisingly, in cells depleted of Sec23, beginning 2 h after induction of a DSB, Pds1 levels were much lower and sometimes undectable (Fig. 3B). Moreover, the Rad53 kinase, whose phosphorylation is indicative of checkpoint activation, was completely dephosphorylated (Fig. 3B). These surprising results suggest that disrupting anterograde traffic by inhibiting COPII vesicles destabilized Pds1 and blocked the maintenance of the DNA damage response.

### DTT treatment after a DSB turns off the DNA damage response

Since COPII vesicles originate in the ER, we considered that degradation of Sec23 and inhibition of COPII vesicles leads to the induction of the ER-stress response, including the unfolded protein response (Walter and Ron, 2011; Jonikas et al. 2009). If this were the case, our data would suggest that the ER-stress response bypasses the DNA damage response, and that both the DNA damage response and the ER-stress response cannot signal simultaneously. To address whether this was indeed the case, we treated cells experiencing a DSB with dithiothreitol (DTT). DTT is a strong reducing agent that blocks the formation of disulfide bonds and leads to the accumulation of unfolded proteins in the ER, activating the unfolded protein response (UPR) and the ER-stress response (24, 40). Cells were grown overnight in YEP-Lac to exponential phase and then treated with 2% galactose to induce a DSB. Cultures were split 2 h after the addition of galactose and one half was treated with 2 mM of DTT. Cells were collected at various time-points to monitor levels of Pds1 and the activation of the DNA damage checkpoint via Rad53 phosphorylation. Interestingly, cells experiencing a DSB and treated with DTT had much lower levels of Pds1 while Rad53 became fully dephosphorylated, compared to cells treated with galactose only, indicative of an inactive DNA damage checkpoint response (Fig. 4A). Moreover, addition of 2 mM DTT 6 h after creating the DSB also released most cells from G2/M arrest after induction of a single HO endonuclease-induced DSB (Fig. 4B). These results suggest that after DTT treatment and activation of the UPR, the DNA damage response is turned off even in the presence of a an irreparable DSB, again suggesting that both the UPR and the DNA damage response cannot be active at the same time.

**Figure 4:**
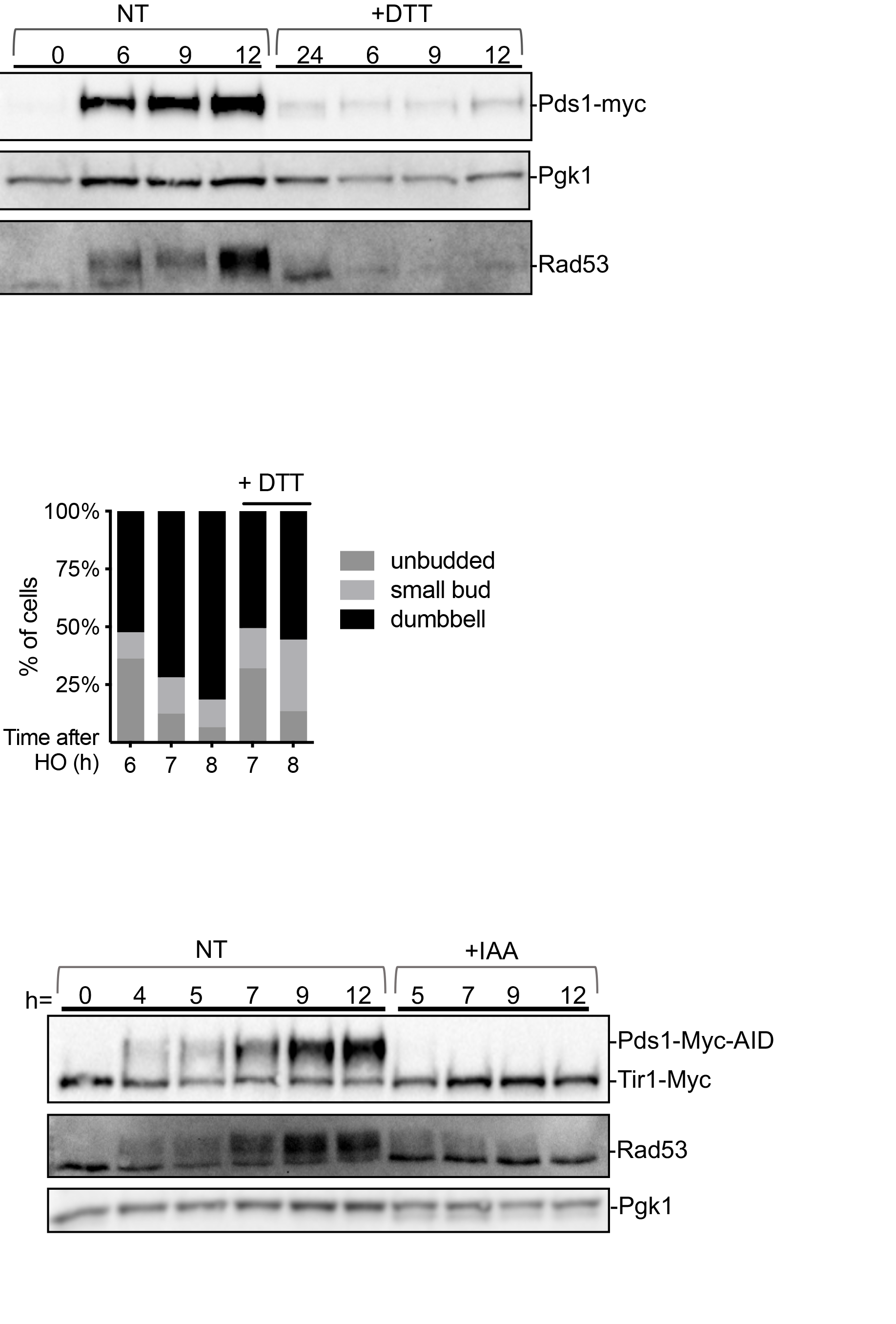
Effect of DTT on the DNA damage checkpoint A) Cells with a galactose-induced HO DSB at t =0 were treated with 2 mM DTT at 6 h and the abundance of Pds1-myc was monitored by Western blot. B) Release of cell cycle arrest when cells with an HO induced DSB were treated with 2mM DTT at 6 h. C) Loss of Rad53 phosphorylation after Pds1-MYC-AID was degraded by IAA treatment.

### Pds1 is required for the maintenance of the DNA damage checkpoint response

Our results indicate that both inhibition of COPII vesicle transport and DTT treatment lead to the degradation of Pds1 and inactivation of the DNA damage checkpoint response. However, we do not know whether these observations reflect a direct role for Pds1 in maintaining the DNA damage response versus simply acting as an effector. To address this question, we tagged Pds1 with AID to enable the degradation of Pds1 after the activation of the DNA damage checkpoint. Cells were grown overnight in YEP-Lac and 2% galactose was added to induce a DSB. 4 h after galactose treatment, cultures were split and 500 uM of auxin was added to degrade Pds1. Pds1 levels and Rad53 phosphorylation were monitored via western blot. In galactose-only treated cells at 4 h, Pds1 protein levels accumulate while Rad53 is phosphorylated, indicative of an active DNA damage checkpoint response (Fig. 4C). In the presence of auxin, Pds1 was rapidly degraded, coinciding with Rad53 dephosphorylation indicating that Pds1 is indeed required for the maintenance of the DNA damage response (Fig. 4C). We suspect that degradation of Pds1 enables Esp1 to cleave sister cohesin rings and allows cell division to proceed, and that progression of the cells from metaphase to anaphase switches off DNA damage signaling.

### Crm1 is an exporter of Pds1 during the DNA damage response

We then asked how Pds1 was exported from the nucleus during DNA damage. Crm1 (Xpo1) is a major exporter in eukaryotic cells (41, 42). Because Crm1 is essential, we again used the AID system and tagged Crm1 with an AID tag tag at the C-terminus to degrade the protein upon IAA addition. Again, cells were grown overnight in YEP-Lac to exponential phase and 1 h prior to galactose induction of a single DSB, we added 500 µM of IAA to promote Crm1 degradation. We monitored Pds1 levels in the absence of Crm1 for a 24 h period in the presence of a single DSB. In the absence of Crm1, we observed higher overall levels of Pds1 during the DNA damage response (Fig. 5D) suggesting that Crm1 is important in displacing Pds1 out of the nucleus during the DNA damage checkpoint response.

**Figure 5:**
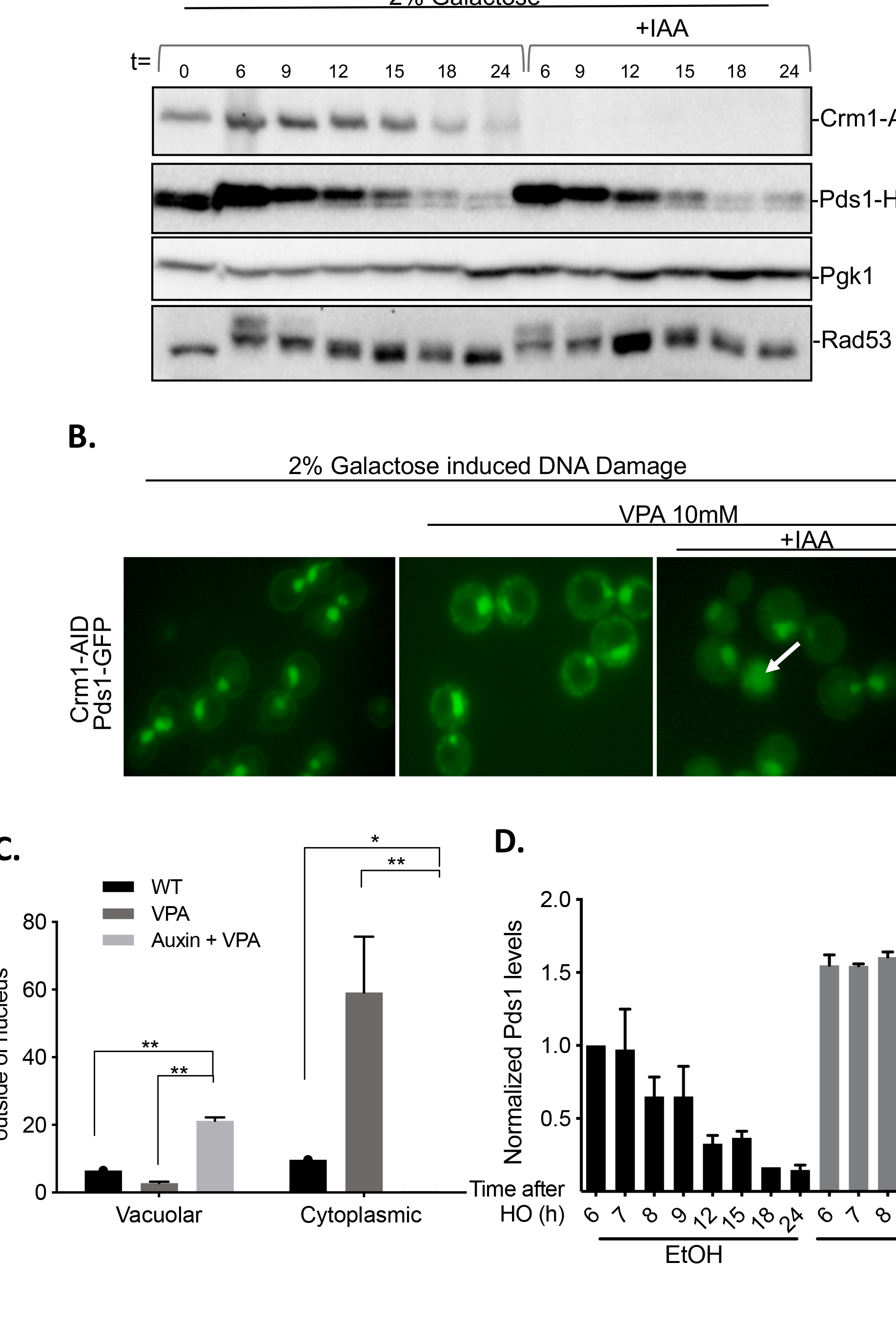
Crm1 is an exporter of Pds1 during the DNA damage response a) JKM179 cells expressing the F-box protein Tir1, Crm1-MYC-AID and Pds1-HA following HO induced DSB b) VPA treated cells in the presence of Crm1 or absence following IAA treatment c) Quantification of Pds1 localization in cells treated with VPA and IAA with VPA. Arrow shows vacuolar localization of Pds1.d) Quantification of Pds1 in cells suffering from HO induced DSB with and without 500 µM of IAA normalized to Pgk1 loading control and Pds1 levels at 6 h in cells treated with vehicle only (EtOH).

To examine further the role of Crm1, we took advantage of the observation that some proteins were dramatically exported from the nucleus after DNA damage after treatment with the histone deacetylase inhibitor, valproic acid (VPA) (43, 44). After inducing a single DSB and allowing cells 6 h to become G2/M arrested we added 10 µM of VPA and Pds1-GFP localization was monitored using fluorescence microscopy. Pds1 localizes to the nucleus during the DNA damage-induced G2/M arrest; however, 30 min after VPA treatment Pds1 rapidly localized to the cytoplasm in puncta-like structures (Fig. 5B). To monitor Pds1 levels after VPA treatment we performed a time-course experiment and quantified Pds1 using western blot analysis. As we have previously shown in this study, Pds1 levels peaked around 6 h, after which levels began to slowly decrease. However, treating cells with VPA significantly reduced Pds1 levels during the DNA damage response (Fig. 5A) In contrast, in Crm1-AID cells treated with IAA prior to start of experiment (Fig. 5A), Pds1 remained in the nucleus for an extended time (Fig. 5B). Very few cells had cytoplasmic Pds1 signal prior to VPA treatment, in both the presence or the absence of Crm1 (Fig. 5C), but in the absence of Crm1 after VPA treatment, Pds1 accumulates in the vacuole, even in the presence of the vacuolar protease Pep4.

## Discussion

This study began with the observation that Pds1-GFP could be seen in the vacuole after DSB damage, in the absence of the vacuolar protease *PEP4* (14). Our initial hypothesis, that Pds1 is targeted to the vacuole in an autophagy dependent manner, proved incorrect, since Pds1 localizes to vacuoles after DSB induction in the absence of core autophagy kinase, Atg1 and regulatory protein Atg13. Instead, we determined that both Pds1 and Esp1 are targeted to the vacuole using the ALP pathway. Although the ALP pathway is named because of the substrate initially described in this pathway (alkaline phosphatase), other proteins such as the vacuolar amino acid transporters Ypq1, Ypq2 and Ypq3 also require components of this pathway (45). We found that inhibition of the ALP pathway causes cells to adapt faster in response to a persistent DSB. This result suggests that proper cell cycle arrest during the DNA damage checkpoint is enhanced by the vacuolar degradation of Pds1. This conclusion is also supported by degrading the nuclear exporter Crm1, since we observe higher levels of Pds1 in its absence. Taken together, these results suggest that proper regulation of Pds1 during the DNA damage response requires nuclear export and traffic to the vacuole. This conclusion is supported by our discovery that inhibition of COPI vesicles significantly affected the localization of Pds1, which now accumulated in the vacuole. These results favor a model where Golgi-accumulated Pds1 gets delivered to the vacuole via AP-3 coated vesicles. This model also supports previous observations that Pds1 is mislocalized in the absence of components of the GARP complex (14). GARP mediates the vesicle transfer from endosomes in the secretory pathway to the trans-Golgi network (46, 47). We suggest that in the absence of GARP, Pds1 is trapped in endosomes and is incapable of re-localizing to the endocytic pathway and back into the nucleus. However, the defects observed in GARP mutants could be rescued either by deleting *PEP4* or by fusing Esp1 with the nuclear localization signal of SV40 to drive Esp1 into the nucleus without relying on its normal chaperone, Pds1 (14).

Many proteins required for protein transport interact with cargo via conserved motifs. Indeed, analysis of Pds1’s protein sequence identifies various protein motifs that are consistent with the export and trafficking pathways in our data. For example, within Pds1 we identify amino acid sequence 112-126 that is consistent with a nuclear export signal (NES) for Crm1. There are also multiple amino acid motifs that have been associated with binding to the mu subunit of the adaptor protein complex AP-3 (48), and while Pds1 has three different Apm3 binding motifs, Esp1 has twenty. Furthermore, both Esp1 and Pds1 have clathrin binding motifs (49, 50) that are consistent with COPII vesicle transport. We have yet to determine whether Pds1 or Esp1 directly interact with vesicle transport proteins to mediate its traffic. Given that Esp1 has a 1630 aa sequence while Pds1 has only 373 aa, Esp1’s larger surface area creates more opportunity for other protein-protein interactions. Pds1 is highly unstructured, but when bound to Esp1 they together form an elongated shape. However a crystal structure of Pds1 bound to Esp1 only resolved residues 258-373 bound to Esp1’s protease site, leaving open the possibility that the C-terminal of Pds1 may have flexibility to interact with other proteins while still inhibiting Esp1’s activity (51).

While looking for proteins that affect Pds1’s localization during the DNA damage response we found that inhibition of COPII vesicles stimulates degradation of Pds1 and dephosphorylation of Rad53. Similar results were observed when cells were treated with DTT, which has been shown to induce the UPR. These results suggest that COPII inhibition during the DNA damage response induces ER stress and that the UPR is dominant over the DNA damage response. Although this crosstalk has not been explored in budding yeast, a similar connection has been made in mammalian cells, where UPR induction after DNA damage prevents the expression of p21 during DNA damage and increases p53’s genotoxic-induced apoptosis (52). Additionally, tunicamycin-induced ER stress has been shown to decrease DNA repair in tumor cells by stimulating Rad51 degradation (53). We have yet to explore the effects of the UPR in DNA repair.

Taken together our results show that Pds1 is highly dynamic in cells during the DNA damage response. Until recently, Pds1 was thought to only localize to the nucleus and only function to regulate Esp1 activity during the cell cycle. Here we show that Pds1 and Esp1 also localize to the vacuole, cytoplasm and Golgi. This multi-compartment localization has been already observed in mammalian cells; in fractionated mammalian cell lysates, securin is found in nuclear, ER, Golgi, and cytoplasmic fractions (54). We have not ruled out the possibility that securin has additional roles outside of the nucleus during the DNA damage response; however, the fact that Esp1 colocalizes with Pds1 in the vacuole during the DNA damage response makes it likely that Pds1 still functions outside of the nucleus to control Esp1’s localization, and to maintain G2/M arrest.

## Materials and Methods

### Media, strains and adaptation assays

All strains used in this study are derivatives of JKM179 (55). All mutant strains were created using one-step PCR homology cassette amplification with lithium acetate transformation procedure (56, 57). AID-tagged strains were constructed as previously described (39). For time-course and imaging experiments, cells were grown overnight in YEP-Lac to early exponential-phase. 2% galactose was added to cultures to induce expression of HO endonuclease. For all AID tagged experiments, 500 µM of 3-Indoleacetic acid (IAA) (Sigma Aldrich) was added to cultures 2 h after galactose addition. Adaptation assays were carried out as previously described (17, 58). Briefly, cells were grown overnight in YEP-Lac and plated in YEP-Lac agarose plates containing 2% galactose. 50 G1 cells were micro-manipulated and monitored for arrest and adaptation, results shown are a replicate of at least three different adaptation study.

### Western Blotting

50 ml of cells from early exponential-phase cultures (approximately 1 x 106 cells/ml) were harvested for each time-point within a time-course experiment and cell pellets were processed using the trichloroacetic acid (TCA) protocol previously described (59). 10%, 8% and 6% Polyacrylamide gels were used in these studies. Blotting was performed using anti-PGK1 (Abcam, ab113687), anti-MYC (Abcam, 9E10), anti-HA (Abcam, HA.C5) and anti-Rad53 (Abcam, ab166859). Blots were imaged using the ChemiDoc Imager (BioRad) and analyzed using the Image Lab (BioRad) software. Protein quantifications were carried out by normalizing protein intensities to Pgk1 loading controls.

### Image Acquisition and Analysis

Cells were grown overnight in YEP-Lac and 2% galactose was added to induce expression of HO. Cells were harvested 6 h after galactose addition and washed in synthetic complete medium before imaging. A Nikon Eclipse E600 microscope was used for imaging. Images were processed and analyzed using the FIJI software. Student’s t-test with a Sidak-Bonferroni correction was performed using GraphPad Prism. The Eukaryotic Linear Motif (ELM) resource (http://elm.eu.org/index.html) was used to analyze the protein sequence of Pds1. The ELM prediction tool scans the protein sequence for motifs corresponding to experimentally validated protein motifs (60).

**Table 1.**
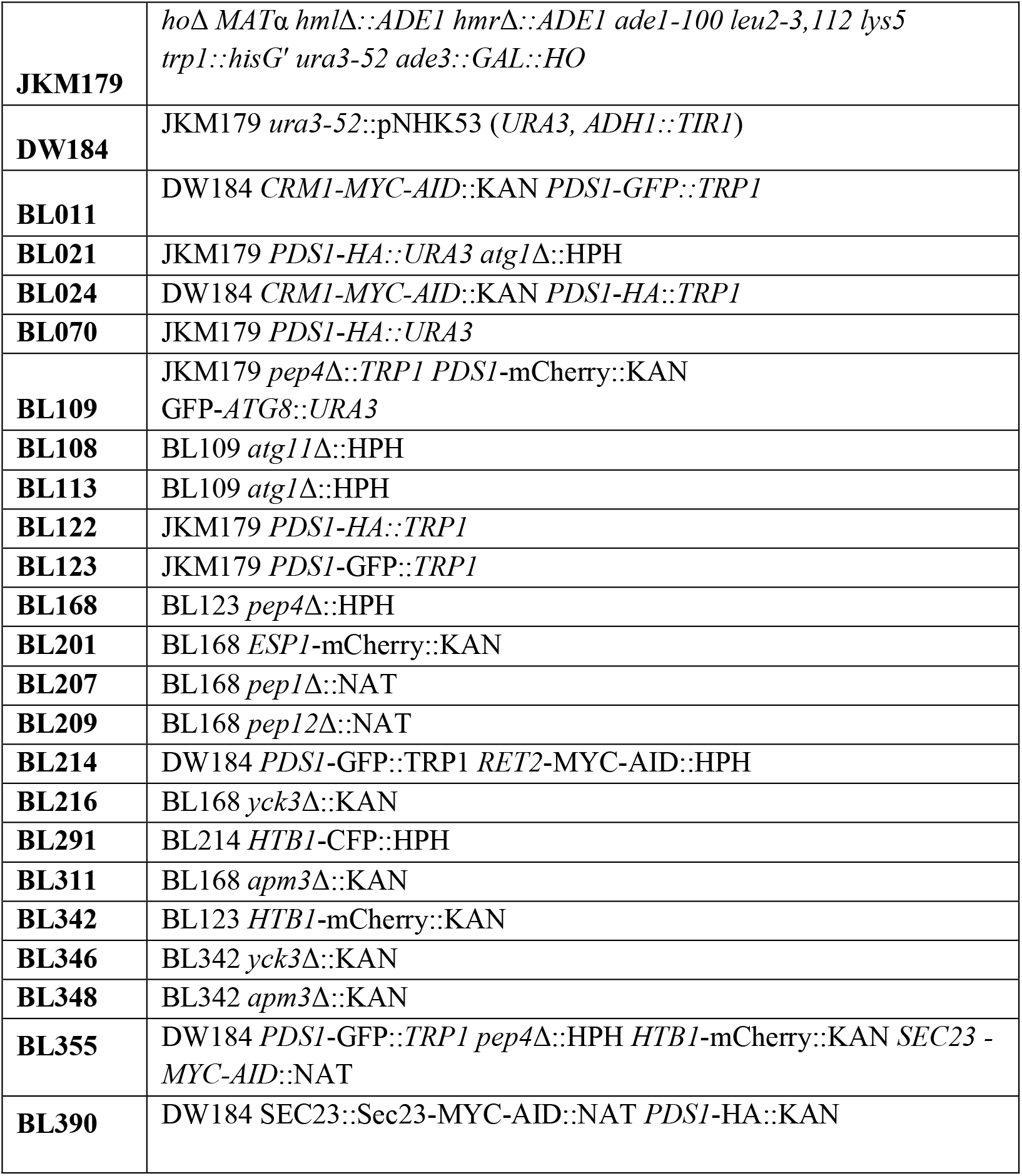
Strains used in this study

## Acknowledgements

We are grateful to Eric Baehrecke and Susan Ferro-Novick for their comments and suggestions on the manuscript. This work was supported by grants GM61766 and R35GM127029. B.L. and D.P.W. were both Trainees of NIH Genetics Training Grant TM32 GM007122. B.L. also received a Research Supplement to Promote Diversity in Health-Related Research for grant R35GM127029.

## Supplemental Information

**Supplemental Figure 1:**
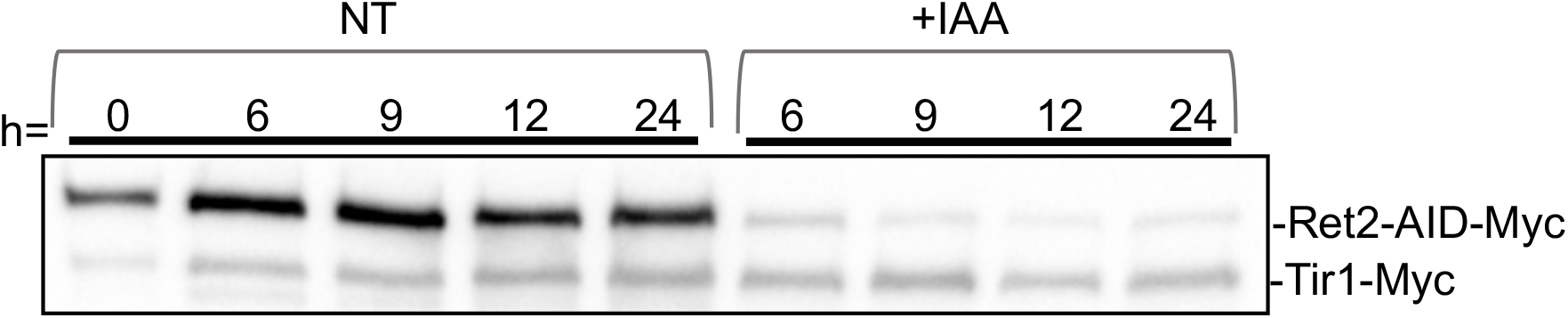
JKM179 cells expressing the F-box protein Tir1 and Ret2-MYC-AID; Ret2 is degraded after IAA treatment, persisting 20 hs after IAA treatment.

